# An *in vitro* assessment of anti-SARS-CoV-2 activity of oral preparations of iodine complexes (RENESSANS)

**DOI:** 10.1101/2020.06.29.171173

**Authors:** Imran Altaf, Muhammad Faisal Nadeem, Nadir Hussain, Muhammad Nawaz, Sohail Raza, Muhammad Asad Ali, Sohail Hasan, Nazish Matti, Muhammad Ashraf, Ihsan Ulla, Sehar Fazal, Saira Rafique, Muhammad Adnan, Nageen Sardar, Tahir Khan, Muhammad Moavia, Sohaib Ashraf, Zarfishan Tahir, Nadia Mukhtar, Tahir Yaqub

## Abstract

Since the emergence of CoVID-19 pandemic in China in late 2019, scientists are striving hard to explore non-toxic, viable anti-SARS-CoV-2 compounds or medicines. We determined *In Vitro* anti-SARS-CoV-2 activity of oral formulations (syrup and capsule) of an Iodine-complex (Renessans). A monolayer of vero cells were exposed to SARS-CoV-2 in the presence and absence of different concentrations (equivalent to 50, 05 and 0.5 μg/ml of I_2_) of Renessans. Anti-SARS-CoV-2 activity of each of the formulation was assessed in the form of cell survival, SARS-CoV-2-specific cytopathic effect (CPE) and genome quantization. With varying concentrations of syrup and capsule, a varying rate of inhibition of CPE, cells survival and virus replication was observed. Compared to 0.5 μg/ml concentration of Renessans syrup, 5 and 50 μg/ml showed comparable results where there was a 100% cell survival, no CPEs and a negligible viral replication (ΔCT= 0.11 and 0.13, respectively). This study indicates that Renessans, containing iodine, may have potential activity against SARS-CoV-2 which needs to be further investigated in human clinical trials.

## Introduction

The SARS-CoV-2 is a novel coronavirus that was first reported in December 2019 in Wuhan, China. The virus was named as 2019nCoV (2019 novel coronavirus) by World Health Organization (1). The International Committee on Taxonomy of Viruses renamed it as Severe Acute Respiratory Syndrome Coronavirus-2 (SARS-CoV-2) on 11 February 2020(2). The infection was termed as Coronavirus Disease (CoVID-19) and its spread forced the World Health Organization to declare it a global pandemic i.e. public health emergency of international concern (PHEIC) (3). As of June 20, 2020, there have been 8,788,791 confirmed cases of CoVID-19 infection, and 463,181 confirmed deaths while 4,648,256 have recovered (4).

Since SARS-CoV-2 infection is a recent emergence in the field of medicine, there is no predefined standard therapeutic course to follow. Most of the treatment regimens revolves around the previous pathophysiological viruses/diseases similar to CoVID-19(5,6). Due to a rapid surge in data regarding CoVID-19, new clinical findings are paving a way forward for more informed decisions in the selection of appropriate therapeutic regimens. Such regimens are focused upon symptomatic relief (e.g. respiratory), amelioration of underlying pathological phenomena (anti-inflammatory) and antiviral effects. With a hope to devise appropriate but definite cure, re-purposing of already available therapeutic options, evidence-based medicine, and traditional therapies are being tested (7) with anecdotal outcomes. For instance, viral fusion inhibitor (arbidol) and protease inhibitors (lopinavir and ritonavir) failed to reduce the negative conversion time of novel coronavirus nucleic acid in pharyngeal swab or improving the symptoms (8). Danoprevir (an anti-HCV drug), on the other hand, has entered phase 4 of clinical trials for treating viral pneumonia in combination with ritonavir (9). Another drug boceprevir has shown to inhibit viral replication during recent experiments (10). These are few examples of projects going-on to discover possible treatment of coronavirus infection, while there are many experiments in clinical trials (11).

Another area is to explore potential micronutrients and vitamins in the treatment of SARS-CoV-2. An extraneous administration of micronutrients are supposed to strengthen the nutritional deficiencies and body’s immune system (12,13). Currently, in the subject matter, many clinical trials are going on to evaluate the efficacy of vitamin D and C in CoVID-19 patients (14,15). A recent study revealed antiviral properties of lithium in preclinical studies for CoVID-19 (17) Though a use of micronutrient is already exemplified by the addition of zinc to chloroquine therapy (16), the direct antiviral potential of micronutrients is still an area wide open for research. Among the micronutrients, Iodine has known antimicrobial properties, and therefore used in topical applications (18). Besides a role for inactivation of enveloped and non-enveloped viruses (19), its use in physical inactivation of SARS-CoV-1, MERS has already been demonstrated (20,21). However, a use of iodine as a systemic intervention is yet debatable for its toxicity (22).

The most common method to increase body iodine concentration is excessive iodine intake through use of iodine supplements (23). Kelp is a dried seaweed rich in vitamins, minerals especially iodine where iodine is complexed with other components. It has traditionally been used for many ailments including weight loss and galactologue (24). An iodine complex formulation has been patented (patent no: 141316, IPO, Pakistan) and registered by MTI, Pakistan (DRAP registration # 505620098). Its clinical trials has successfully been conducted where it was found to be highly effective in the treatment of oligomenorrhea and polycystic fibrosis (25,26). In addition to that, antiviral efficacy of the studied drug has also been reported for clinical trial against Hepatitis C Virus (HCV). Indeed, in combination with traditional therapy, iodine complex has been associated with excellent antiviral response in chronic HCV patients (27). Antiviral activity of said complex (Renessans) has also been tested for avian influenza virus where it showed an inhibition of cytopathic effects (28).

With this background, considering previous potential of study drug, we investigated antiviral potential of Iodine complex simply because it is readily available for public use, and has a proven antiviral efficacy of iodine complexes.

## MATERIAL AND METHODS

### Drugs

RENESSANS capsule (containing 200mg iodine) and syrup (containing 10mg/ml iodine) were prepared and provided by MTI Medical Pvt. Ltd. for this study.

#### Revival and establishment of VERO cell line monolayer

Vero cells were obtained from UVAS WTO-Quality Control Lab, University of Veterinary Animal Sciences Lahore, Pakistan. Cells were revived using DMEM cell culture media using 10% Fetal Bovine serum. Monolayer was established and maintained in 25cm^2^ roux flasks. The flasks were observed for 48 hours for any kind of bacterial or fungal contamination. The flasks with 80% monolayer were selected for viral replication and antiviral activity of drugs.

#### Virus cultivation and isolation

SARS-CoV-2 (nCOV-1-20-IoM) was isolated previously from a clinical sample in BSL- 3 laboratory located in the Department of Microbiology, University of Veterinary and Animal Sciences, Lahore. The isolate was identified using commercially available real-time PCR kit (Sansure BioTech, China) as per manufacturer’s instruction. Cell flasks with 80% monolayer were used and infected with SARS-CoV-2 isolate. DMEM cell culture media containing 1% fetal bovine serum was used for infection. After 48-72 hours incubation, cytopathic effects were observed and virus replication was confirmed by real time RT-PCR.

#### Preparation of drug concentrations

Different concentrations of drugs were prepared and reconstituted in same media used for Vero cell. Firstly the cytotoxicity profile of drugs was checked and non-cytotoxic concentrations were selected for the evaluation of antiviral profile in-vitro cell culture. The concentrations selected for study were 100 μg/ml, 10 μg/ml and 1μg/ml.

#### Cytotoxicity Assay

Different drug concentrations that were selected for antiviral activity were mixed into cell culture media and added into confluent VERO cell line with 80% monolayer. Control flasks were also kept by adding 10% DMSO in them. Flasks were then incubated at 37°C in 5% Co_2_.

#### In-vitro Antiviral activity

Different concentrations of the drugs were prepared and reconstituted in DMEM. The concentrations were evaluated for the cytotoxicity profile in vitro cell culture. The noncyototoxic concentrations (100, 10 and 1 μg/ml of I_2_) were selected for the evaluation of antiviral profile in vitro cell culture against SARS CoV-2. One milliliter (1ml) of respective concentrations of drug was mixed DMEM (1 ml). This mixture was then added on vero cell line monolayer and incubated for 30 minutes. After 30 minutes, drug mixture was removed and cells were washed with media. One ml of each respective concentrations of drug (100, 10 & 1 ug/ml) were mixed with I ml of viral suspension in DMEM (CT value was adjusted to 20 by quantitative real time PCR) and was incubated for 30 minute at room temperature. Respective drug-virus mixtures were added in separate flasks containing vero cell line for the evaluation of in vitro antiviral activity of the selected concentrations of drugs. The flasks were incubated at 37°C in 5% Co_2_ for 72 hours and then checked for cytopathic effects, and viral load which was estimated in terms of CT values measured by RT-PCR. Positive control cell flask was inoculated with equal amount of SARS-CoV-2 while and no drug was added.

### Genome Extraction and quantitative RT PCR

The RNAs were extracted using the Systasq SuperExtract viral RNA purification Kit (Systaaq Diagonostic Products, USA) as per manufacturer’s instruction.SARS-CoV-2 was detected using real time RT-PCR [Sansure Biotech Novel Coronavirus (2019-nCoV) Nucleic Acid Diagnostic Kit (Sansure Biotech, China)] as per manufacturer’s instructions. The kit has a potential to simultaneously detect both ORF1ab and N genes corresponding to SARS-CoV-2. Virus load was estimated by measuring the change in Ct values of among drug-treated and drug-untreated or normal cells.

## Results

The VERO cells were exposed to SARS-CoV-2 with and without different concentration of Renessans capsule and syrup. Effect of renessans capsule and syrup on the growth of virus on vero cells monolayer is given in Table 1. Positive control flasks had typical SARS-CoV-2 cytopathetic effects within 48 hours where no cell was found survived. Negative control had no CPEs and 100% cells were survived (Figure 1). A lack of CPEs was observed on vero cells exposed to higher concentrations of Renessans syrup (50 μg/ml and 5 μg/ml of I_2_) with an excellent (100%) cell survival. Vero cells exposed to a lower concentration of Renessans syrup (equivalent to 0.5 μg/ml I_2_), and all concentrations of Renessans capsule (equivalent of 50, 5.0 and 0.5 μg/ml of I_2_) showed 90% survival, after SARS-CoV-2 exposure and 10% (Table 1, Figure 1).

**Table 1:** Effect of RENESSANCE syrup and capsule on cytopathic effect of SARS CoV-2 on in vitro VERO Cells culture system.

**culture**
| Formulation | Conc.<br>(µg/ml) of<br>drug | Cell Survival<br>% | % CPE |
| --- | --- | --- | --- |
| Renessance<br>Syrup | 50 | 100 | 0 |
|  | 5 | 100 | 0 |
|  | 0.5 | 90 | 10 |
| Renessance<br>Capsule | 50 | 90 | 10 |
|  | 5 | 90 | 10 |
|  | 0.5 | 90 | 10 |
| SARS CoV-2<br>Control (PC) | None | 0 | 100 |
| Cell control<br>(NC) | None | 100 | - |

**Figure 1:**
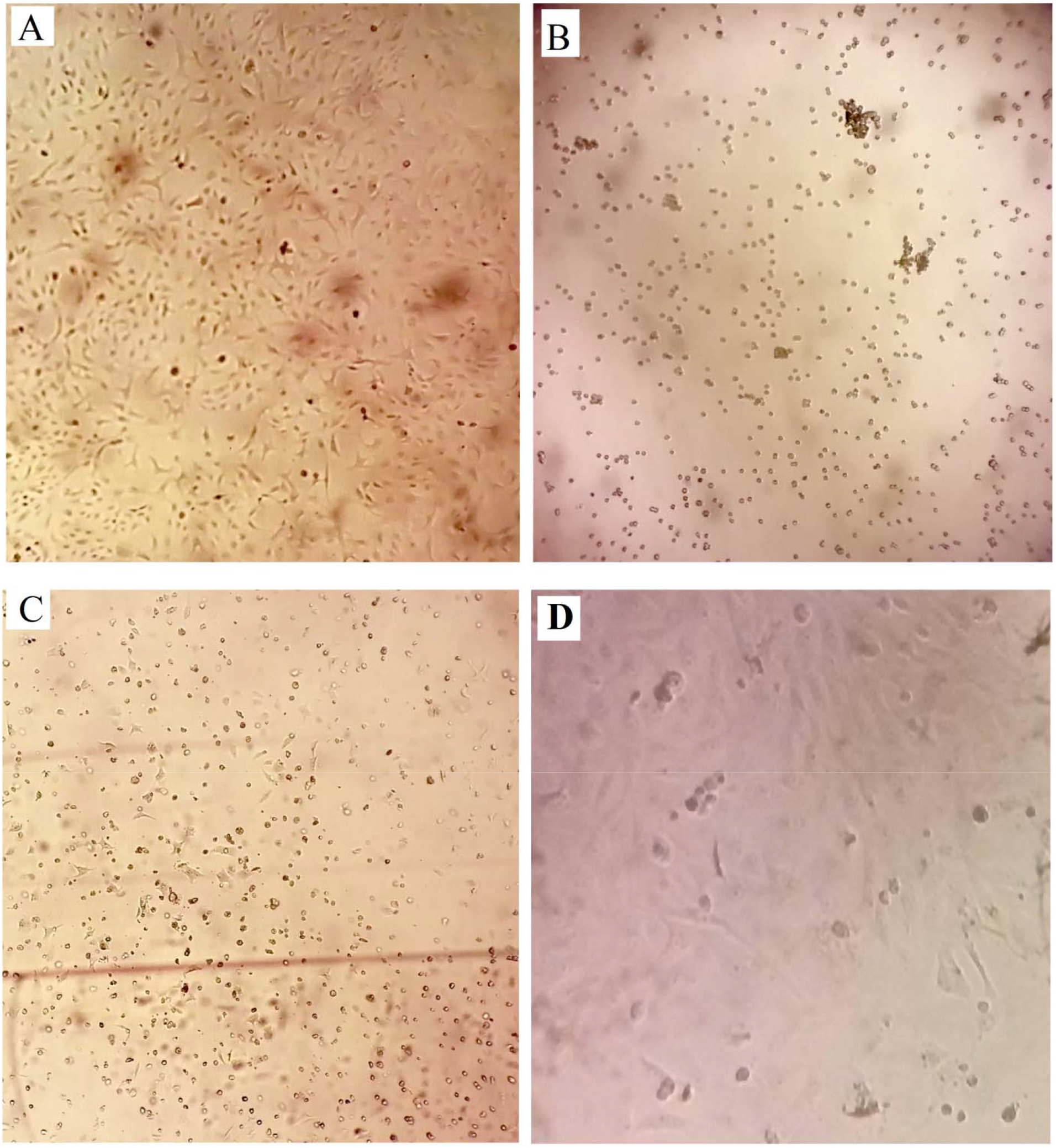
Effect of RENESSANS on cytopathetic effect of SARS CoV-2 on VERO Cells. A: VERO Cells showing no CPEs and 100% cell survival; B) VERO Cells with 100% CPEs and no cell survival after exposure to SARS-CoV-2; C) Vero cells 90% CPEs and 10% Cell survival after exposure to SARS-CoV-2; D) Vero cell showing 10% CPEs after exposure to SARS-CoV-2

The SARS-CoV-2 was detected from all the treated and untreated vero cells by Real time RT-PCR (Table 2). Detection cycle threshold (Ct) was used to estimate the viral load and ΔCt was used as an indicator of viral replication on vero cells. The Ct of the inoculated flask, immediately after virus inoculation, was found to be 19.5. In absence of any drug, Ct value was reduced to 12.4 (Δ**Ct=7.6**), indicating a significant replication of SARS CoV-2. A higher concentration of Renessans syrup showed minimum ΔCt (0.11 and 0.13, respectively), while ΔCt for higher concentration of Renessans syrup was found to be 1.46 which is a clear indicator for potential inhibition of virus and a subsequent decrease in genome copies corresponding to SARS-CoV-2. All the concentrations of RENESSANS capsule showed similar effect on ΔCt (1.4) of SARS-CoV-2 detection by real time RT-PCR.

**Table 2:**
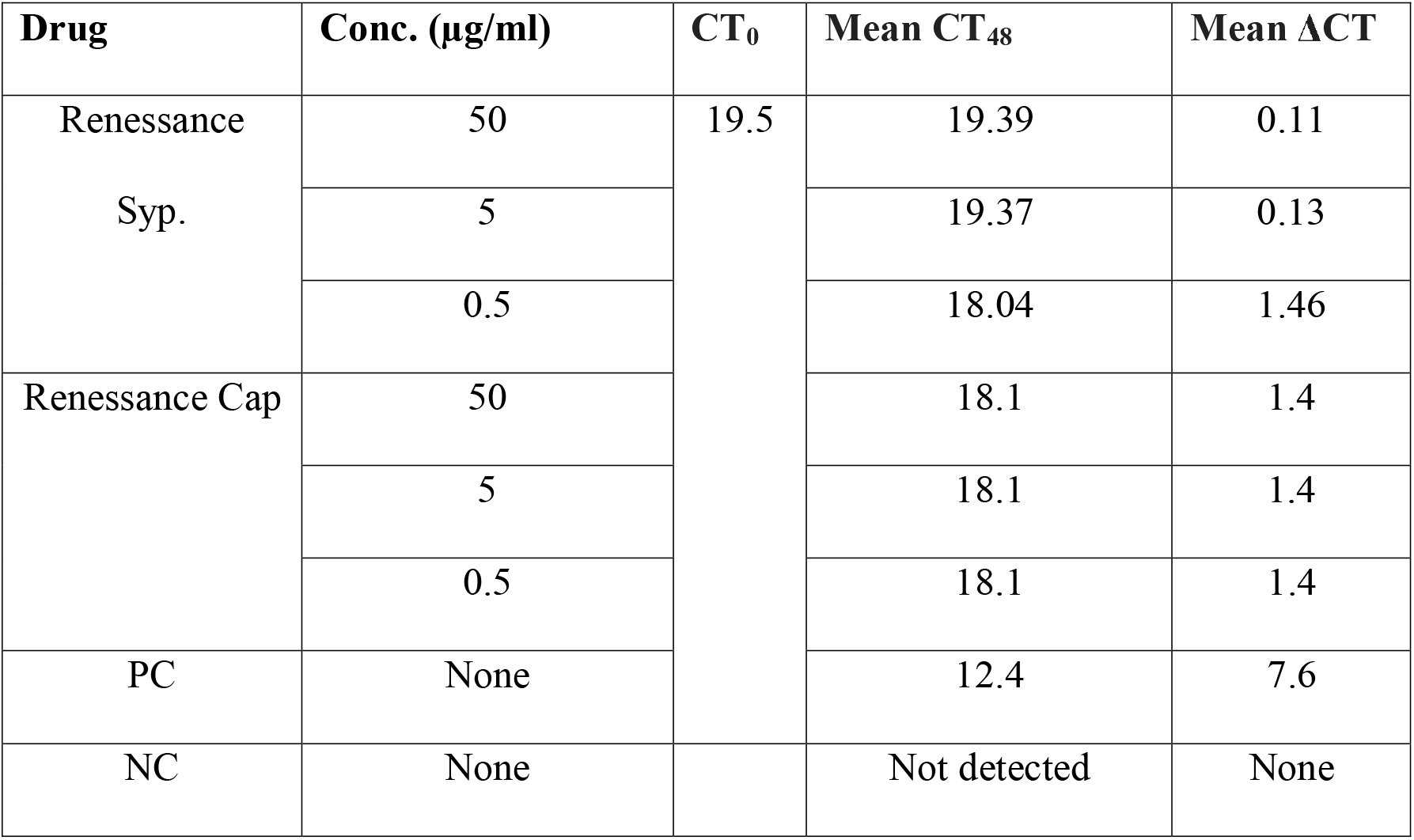
Effect of RENESSANCE syrup and capsule SARS CoV-2 growth in VERO Cells.

| Drug | Conc. (µg/ml) | CT<sub>0</sub> | Mean CT<sub>48</sub> | Mean ΔCT |
| --- | --- | --- | --- | --- |
| Renessance<br>Syp. | 50 | 19.5 | 19.39 | 0.11 |
|  | 5 |  | 19.37 | 0.13 |
|  | 0.5 |  | 18.04 | 1.46 |
| Renessance Cap | 50 |  | 18.1 | 1.4 |
|  | 5 |  | 18.1 | 1.4 |
|  | 0.5 |  | 18.1 | 1.4 |
| PC | None |  | 12.4 | 7.6 |
| NC | None |  | Not detected | None |

## DISCUSSION

COVID-19 pandemic is caused by SARS-CoV-2 COVID-19 pandemic is causing is the most significant threat to the lives on earth today. It is showing no signs of slowing down. On one hand, the world is racing to find the cure against this newly emerged virus through developing vaccines and antivirals. Currently, more than 150 vaccines are in the different phases of development. Most of the infectious disease experts believe that it may require at least 18 months for the first vaccine to be available in the market (29). Indeed, antiviral drug is considered to be the most probable and urgent cure for the COVID-19. However, new development of antiviral against SARS-CoV-2 may require years that initiates the potential repurposing the existing approved antivirals/antimicrobials against SARS-CoV-2. In this context, ivermectin (30), Hydroxychloroquin (31), Arbidol (32) and many others have been tested against SARS-CoV-2. WHO drops HCQ from clinical trials after available data indicated that the drug has no effect against COVID-19 (33). Ivermectin exposure to cells exhibited significant reduction in virus titers as compare to control cells. The trials for the clinical use of Ivermectin against COVID-19 are in progress.

In this study, we evaluated the anti SARS-CoV-2 activity of iodine complex (Renessans) that has already been approved for human use. Iodine has a history of use in tropical applications and exhibited antiviral activity against SARS-CoV, MERS, avian influenza virus and HCV. We have used non toxic concentrations (50μg/ml, 5μg/ml, 0.5μg/ml) of Renessans. At these concentrations, Renessans exhibited strong antiviral activity against SARS-CoV-2 with no/few CPE were observed in drug treated cells as compare to control cells. In vitro results exhibited that Renessans in the form of Syrup able to complete inhibition of CPE (0%) formation at 50μg/ml, 5μg/ml as compare to tablet, which able to inhibit 90% CPE formation at same concentrations. This may predict the use of Renessans in the form of syrup as compare to capsule form. This indicated that Renessans in the syrup formulation has better absorption in the cells than capsule form. In line with the cell morphological analysis, qRT-PCR data revealed that the virus replication was greatly inhibited in the drug treated cells as compare to control cells. In a recent study, the antiviral activity of CupriDyne, an iodine complex disinfectant solution was evaluated against SARS-CoV-2. This iodine solution able to inactivate virus in time dependent manner, reducing the virus titers by 99% and reducing the virus titers below detection limit after 60 min (34). Similarly, iodine complex had exhibited a virucidal activity against MERS virus, virus inactivation of ≥99.99% within 15 seconds of application. Moreover, iodine product had reduced the SARS-CoV infectivity to undetectable levels in 2 minutes of exposure in Vero infected Cells (35). Collectively, the previous data confirms our finding of iodine complexes, which have showed strong antiviral activity against SARS-CoV2 and members of this family. Moreover, iodine has also exhibited its antiviral potential against other viruses like in human and avian influenza virus, iodine able to inhibit the influenza A viruses infection up to 97.5 % in MDCK infected cells (36), adenoviral conjunctivitis (37), and Modified vaccinia virus Ankara (38). Based on previous literature addressing the mechanism involved in the activity of iodine against SARS, it is more likely that iodine makes the structural changes on the viral coat through attack on histidine and tyrosine residues (34). Thus, it is likely that inhibition of SARS-CoV-2 infection at the entry level by blocking the viral attachment to the cell. Indeed, this seems to be a general mechanism underlying the inhibitory effect of iodine on other viruses, including human and avian influenza viruses. Limitation of the study is that we only used CT values for estimation of virus titers; further experiments are needed where TCID_50_ should be determined as well and we are working on that.

This study indicates that RENESSANS (iodine containing oral formulation), may have potential activity against SARS-CoV-2 which needs to be further investigated in human clinical trials.

